# Stimulating stopping? Investigating the effects of tDCS over the inferior frontal gyri and visual cortices

**DOI:** 10.1101/723296

**Authors:** Christina N. Thunberg, Mari S. Messel, Liisa Raud, René J. Huster

**Author notes:** **Correspondence:** René J. Huster, Department of Psychology, University of Oslo, Norway.

## Abstract

The ability to cancel an already initiated response is central to flexible behavior. While several different behavioral and neural markers have been suggested to quantify the latency of the stopping process, it remains unclear if they quantify the stopping process itself, or other supporting mechanisms such as visual and/or attentional processing. The present study sought to investigate the contributions of inhibitory and sensory processes to stopping latency markers by combining transcranial direct current stimulation (tDCS), electroencephalography (EEG) and electromyography (EMG) recordings in a within-subject design. Active and sham tDCS were applied over the inferior frontal gyri (IFG) and visual cortices (VC), combined with both online and offline EEG and EMG recordings. We found evidence that neither of the active tDCS condition affected stopping latencies relative to sham stimulation. Our results challenge previous findings suggesting that anodal tDCS over the IFG can reduce stopping latency and demonstrates the necessity of adequate control conditions in tDCS research. Additionally, while the different putative markers of stopping latency showed generally positive correlations with each other, they also showed substantial variation in the estimated latency of inhibition, making it unlikely that they all capture the same construct exclusively.

## Introduction

Response inhibition, i.e. the ability to cancel a prepotent or ongoing response, is considered essential for adaptive behavior. It is thought to rely on the right inferior frontal gyrus^1,2^ and is often investigated using the stop-signal task (SST). Here, participants are instructed to quickly respond to the appearance of a go-stimulus. In a minority of trials, the go-stimulus is followed by a stop-signal with a variable delay (stop-signal delay; SSD), prompting the participant to cancel their initiated response. Consequently, successful response inhibition is characterized by the lack of observable behavior, making the inhibitory process difficult to investigate.

Several indirect measures have been suggested to quantify the timing of the inhibitory process. The most common measure derived from the SST is the stop-signal reaction time (SSRT). The SSRT is defined as the covert latency of the stopping process^3^, and is commonly calculated as a difference measure between some summary statistic of the reaction times in go trials and the stop-signal delay. At the neural level, the P3 event-related potential (ERP) was early on suggested as a marker of successful inhibition^4^. While the P3 amplitude scales with factors like inhibitory load and incentives for stopping^5,6^, the evidence for an association between P3 amplitudes and SSRTs is equivocal^7–9^. Furthermore, P3 amplitude differences, both between groups and between experimental conditions, have repeatedly been reported without concurrent differences in SSRTs^10–12^. Recently, it has been proposed to shift the focus towards temporal aspects of the P3. In fact, studies have shown that both the peak and onset latency of the P3 might correlate with the SSRT^13–16^, suggesting that P3 latency measures could be potential neural markers of inhibitory capability. In addition to this, studies have found that the effects of inhibitory control can be measured by electromyographic (EMG) activity at response effector muscles in successful stop trials^4,17,18^. While the onset of this so-called partial response EMG (prEMG) reflects the response to the preceding go stimuli, the point for the prEMG decline (i.e. peak latency) may indicate the timing of inhibition reaching the periphery and could thus be another potential marker for the stopping latency.

While a great deal of research has focused on quantifying the stopping latency, it has been argued that the use of inhibition as a general construct is not enough to properly explain SST performance^19–21^. In fact, the computational model underlying SSRT estimation makes no assumptions about the processes leading up to the estimated stopping latency. This suggests that potential stopping markers could be sensitive to variations in processing stages preceding inhibition, a notion which has been supported by recent computational work^22,23^. In fact, several studies show that sensory, attentional and perceptual processes all play into inhibitory performance in the SST^24–27^.

A promising avenue for delineating the relative impact of sensory processing and inhibitory control to the latency of the different potential inhibitory markers is transcranial direct current stimulation (tDCS). TDCS has the potential to modulate neural activity, with anodal and cathodal stimulation being associated with increased and decreased excitability in cortical areas, respectively^28^. Therefore, tDCS can be a powerful tool to make causal inferences about the role of the regions and processes involved in SST performance. Several studies have found decreased SSRTs following anodal stimulation of the IFG^29–34^, suggesting that this method can indeed alter inhibition latencies. However, both increased and decreased reaction times in go trials have also been reported^10,31,35^, indicating that anodal IFG stimulation could potentially influence strategic adjustments of task performance rather than an inhibitory process per se. Furthermore, investigations of concurrent electrophysiological modulations are sparse, and no one has investigated the effects of anodal IFG stimulation on inhibition at the effector level.

In this study, we explored the relationship between the different potential stopping latency measures, i.e. the SSRT, the prEMG, and the peak and onset latency of the P3 potential. We also investigated how these different markers were modulated by tDCS. We used a repeated measures design with three separate tDCS sessions for each participant and compared anodal stimulation of the inferior frontal gyri (IFG) and visual cortices (VC) to sham stimulation. We used a multimodal approach and combined the tDCS with electroencephalography (EEG) and electromyography (EMG) recordings prior to, during, and after stimulation. Based on previous literature, we expected to find decreased SSRTs following anodal IFG stimulation. Additionally, if the SSRT, the P3 onset latency and the prEMG all index the same inhibitory process, we would expect to see decreased SSRTs accompanied by earlier P3 onset latencies as well as earlier prEMG latencies. Furthermore, if anodal tDCS over the visual cortices can facilitate sensory processing, which presumably contributes to the stopping latency, we would expect to see similar decreases in SSRTs, P3 latencies and prEMG latencies following VC stimulation, accompanied by more global decreases in reaction times in go trials.

## Results

To examine the effect of tDCS over regions associated with inhibitory control and sensory processing, we applied 20 minutes of 2mA bilateral anodal tDCS over the IFG, VC as well as sham stimulation. Electrode positions were chosen in accordance with current flow simulations for the specified target areas^36^. All behavioral, EEG, and EMG measures were analyzed used 3 × 3 repeated measures ANOVA (rmANOVA) with the factors **time** (pre-, peri-, post-stimulation) and **condition** (IFG, VC, SHAM). The analyses were performed within a Bayesian framework using the JASP software^37^, and Bayes factors (BF_10_) are reported. BFs are a measure of strength of evidence in favor of a hypothesis^38^. In the case of BF_10_, values below 1 support the null hypothesis, and values larger than 1 supports the alternative hypothesis. It has further been suggested that the strength of evidence can be assessed as follows: values between 0.33-1 or 1-3 are considered inconclusive, only providing anecdotal evidence. Values between 0.10-0.33 or 3-10 are considered moderate, and values smaller than 0.10 or larger than 10 are considered as strong evidence for H0 or H1, respectively^39,40^.

### Behavioral results

Across all measurements, i.e. across pre-, peri- and post-stimulation measurements for all tDCS conditions, average go reaction times (goRT), SSRTs and stop accuracies were 500 ms, 233 ms and 49.5 %, respectively, and thus in line with previous studies using a visual SST in a sample of healthy young adults. Furthermore, goRTs were longer than unsuccessful stop RTs (USRTs) for all participants in all measurements (all BF_10_ > 552 000), thus meeting the assumptions of the horse-race model. A full overview of all behavioral measures can be seen in Table 1.

**Table 1.** Behavioral results and EEG and EMG latencies (N = 18)

|  | IFG |  |  | VC |  |  | SHAM |  |  |
| --- | --- | --- | --- | --- | --- | --- | --- | --- | --- |
|  | pre | peri | post | pre | peri | post | pre | peri | post |
| **GoRT (ms)** | 497 | 484 | 490 | 521 | 511 | 507 | 514 | 491 | 486 |
|  | (61) | (67) | (63) | (59) | (57) | (49) | (70) | (67) | (54) |
| **SSRT (ms)** | 233 | 235 | 238 | 229 | 229 | 235 | 228 | 242 | 231 |
|  | (24) | (20) | (24) | (28) | (17) | (25) | (22) | (22) | (24) |
| **USRT (ms)** | 447 | 445 | 447 | 473 | 472 | 465 | 473 | 449 | 445 |
|  | (58) | (64) | (57) | (54) | (59) | (53) | (62) | (58) | (43) |
| **SSD (ms)** | 264 | 249 | 252 | 291 | 286 | 275 | 288 | 254 | 259 |
|  | (68) | (81) | (74) | (63) | (69) | (64) | (77) | (74) | (64) |
| **Stop-accuracy (%)** | 49.23 | 48.66 | 49.18 | 50.21 | 50.05 | 50.00 | 49.95 | 48.71 | 49.23 |
|  | (3.50) | (3.18) | (3.24) | (1.64) | (1.40) | (1.56) | (1.63) | (5.33) | (2.04) |
| **Go-accuracy (%)** | 98.83 | 98.39 | 98.51 | 98.52 | 98.62 | 98.55 | 98.73 | 99.06 | 98.57 |
|  | (0.72) | (1.65) | (1.93) | (2.39) | (2.62) | (1.89) | (2.02) | (1.31) | (1.57) |
| **P3 peak (ms)** | 369 | 382 | 384 | 363 | 371 | 363 | 371 | 373 | 385 |
|  | (40) | (28) | (34) | (39) | (38) | (42) | (45) | (47) | (31) |
| **P3 onset (ms)** | 302 | 316 | 303 | 295 | 301 | 303 | 301 | 300 | 295 |
|  | (28) | (35) | (28) | (31) | (32) | (32) | (29) | (35) | (30) |
| **prEMG peak (ms)** | 159 | 161 | 168 | 160 | 157 | 154 | 155 | 166 | 161 |
|  | (23) | (27) | (26) | (15) | (20) | (27) | (23) | (23) | (25) |

Estimates represent mean values, with standard deviations represented in parentheses.
Abbreviations: GoRT – go reaction time, SSRT – stop-signal reaction time, USRT – unsuccessful stop reaction time, SSD – stop signal delay, prEMG – partial response EMG

Neither tDCS condition was associated with changes in SSRTs, as indicated by moderate evidence against a **time x condition** interaction (BF_10_ = 0.321). Additionally, SSRTs did not change over the course of a session (BF_10_ = 0.249 for the main effect of **time**), nor did it differ across conditions (BF_10_ = 0.120 for the main effect of **condition**).

TDCS did not affect goRTs, as shown by strong evidence against the **time x condition** interaction (BF_10_ = 0.088). We found anecdotal evidence for a main effect of **time** (BF_10_ = 1.297) and moderate evidence for a main effect of **condition** (BF_10_ = 8.838). Post hoc tests for the **condition** factor suggested that reaction times were longer in the VC session, with strong evidence for a difference between VC and IFG (BF_10_ = 11.703, *M*_diff_ = 23 ms), anecdotal evidence for a difference between VC and SHAM (BF_10_ = 2.683, *M*_diff_ = 16 ms), and moderate evidence against a difference between IFG and SHAM (BF_10_ = 0.239, *M*_diff_ = 7 ms).

In addition, there was no stimulation-specific effects on SSDs, as indicated by strong evidence against the **time x condition** interaction (BF_10_ = 0.094). We found anecdotal evidence against a main effect of **time** (BF_10_ = 0.932) and strong evidence for a main effect of **condition** (BF_10_ = 13.283). Post hoc tests for the **condition** factor suggested that SSDs were longer in the VC session, with strong evidence for a difference between VC and IFG (BF_10_ = 18.163, *M*_diff_ = 28 ms), anecdotal evidence for a difference between VC and SHAM (BF_10_ = 1.260, *M*_diff_ = 16 ms) and anecdotal evidence against a difference between IFG and SHAM (BF_10_ = 0.374, *M*_diff_ = 12 ms).

### EEG results

Across all measurements, the stop-P3 peaked at 374 ms, with an average onset latency of 302 ms. ERPs can be seen in Figure 1a, and P3 latency estimates can be seen in Table 1.

**Figure 1.**
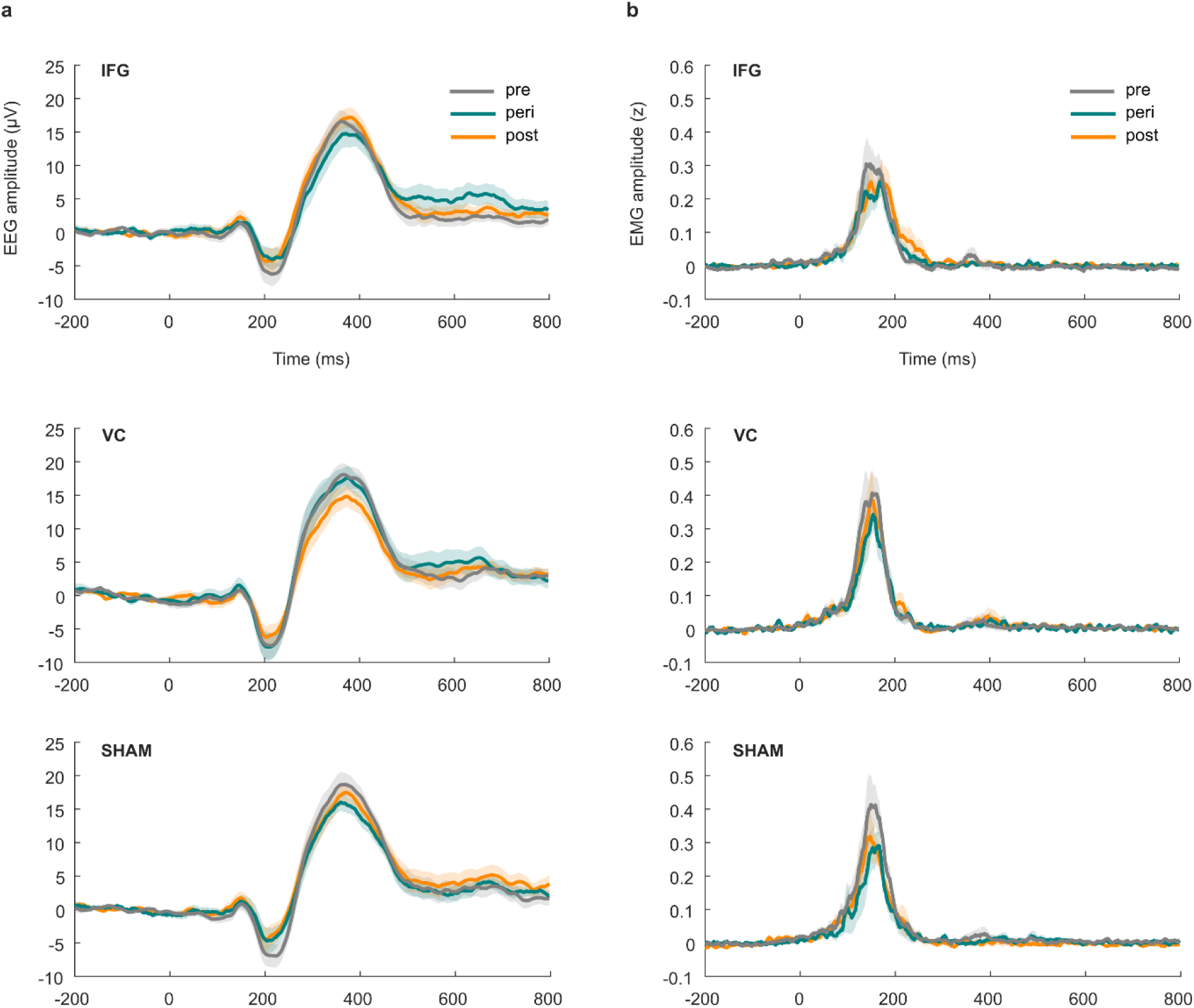
**a.** EEG activity at Cz in successful stop trials, time-locked to stop-stimulus onset. **b.** PrEMG activity in successful stop trials, time-locked to stop-stimulus onset and depicted as z-scored signal strength relative to the pre-go-stimulus-baseline. Shaded areas represent *SEM*.

TDCS did not modulate P3 peak latencies, as shown by moderate evidence against a main effect of **time** (BF_10_ = 0.228), anecdotal evidence against a main effect of **condition** (BF_10_ = 0.811), and moderate evidence against a **time x condition** interaction (BF_10_ = 0.108). However, the interaction term violated the sphericity assumption (Greenhouse-Geisser ε = .61), and as correction procedures are not well developed within the Bayesian framework, some caution is required in in the interpretation of the interaction effect. For the P3 onset latency, there was anecdotal evidence against a main effect of **time** (BF_10_ = 0.514) and moderate evidence for a main effect of **condition** (BF_10_ = 3.353). Post hoc comparisons for the condition factor showed moderate evidence against a difference between VC and SHAM (BF_10_ = 0.159, *M*_diff_ = 1 ms), and anecdotal evidence for a difference between the IFG- and VC-condition (BF_10_ = 1.477, *M*_diff_ = 7 ms) as well as between the IFG-and SHAM-condition (BF_10_ = 2.559, *M*_diff_ = 8 ms), both in the direction of slightly later onset latencies in the IFG condition. The **time x condition** term showed anecdotal evidence against the interaction (BF_10_ = 0.693). Once again, though, a sphericity violation (Greenhouse-Geisser ε = .56) warrants some caution in interpreting the results.

### EMG results

PrEMG activity could be detected in roughly 30 % of successful stop trials, with an average peak latency of 160 ms across all measurements. PrEMG time courses can be seen in Figure 1b, and prEMG peak latencies can be found in Table 1.

The number of detected responses did not differ between measurements, as shown by strong evidence against a main effect of **time** (BF_10_ = 0.091), moderate evidence against a main effect of **condition** (BF_10_ = 0.204), and moderate evidence against the **time x condition** interaction (BF_10_ = 0.219). As for the latency of the prEMG, we found strong evidence against a main effect of **time** (BF_10_ = 0.095), moderate evidence against a main effect of **condition** (BF_10_ = 0.236) and moderate evidence against a **time x condition** interaction (BF_10_ = 0.328).

### Brain-behavior correlations

To get an indication of the stability of the direction of the obtained correlations, we tested the correlation coefficients against the null hypothesis. Average *r* values (*M*_*r*_) and BF_10_ for one-sample t-tests are reported, and an overview of all correlation coefficients between the different measures can be seen in Table 2.

**Table 2.**
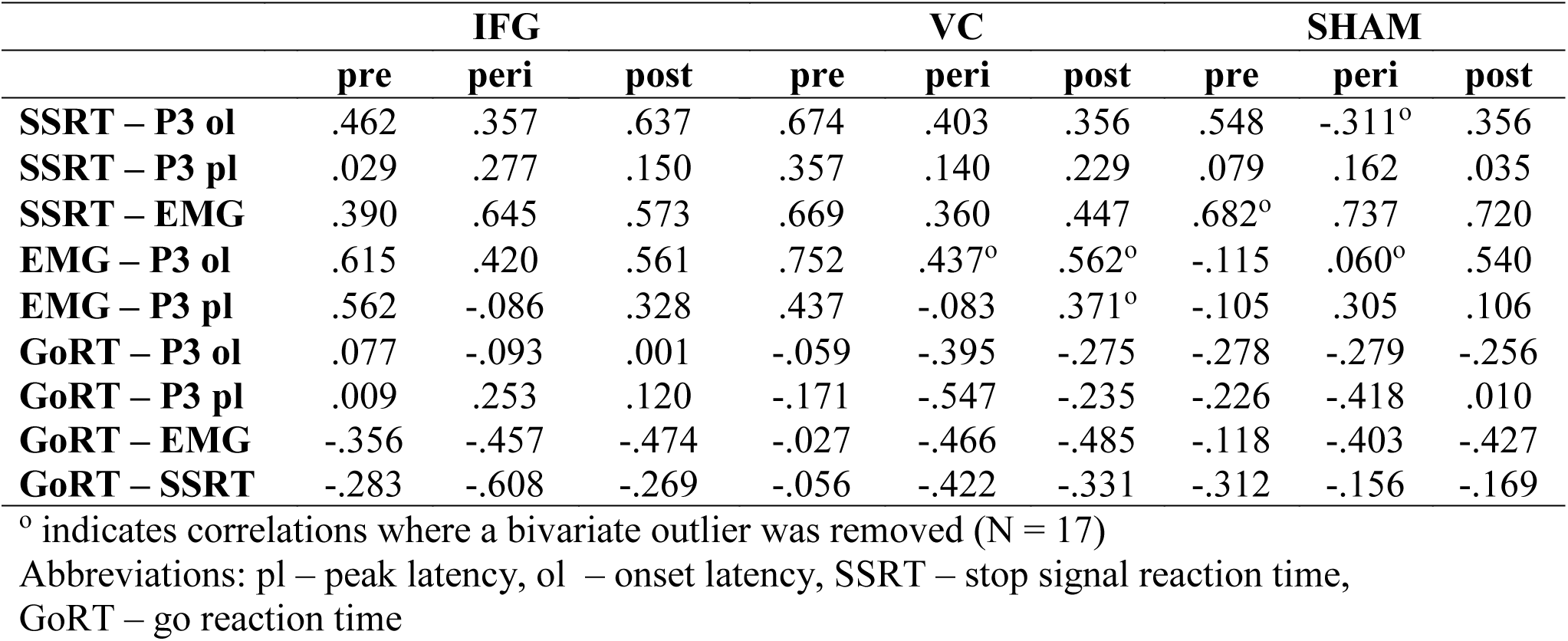
Pearsons *r* between different inhibition measures and reaction times (N = 18)

|  | IFG |  |  | VC |  |  | SHAM |  |  |
| --- | --- | --- | --- | --- | --- | --- | --- | --- | --- |
|  | pre | peri | post | pre | peri | post | pre | peri | post |
| **SSRT – P3 ol** | .462 | .357 | .637 | .674 | .403 | .356 | .548 | -.311<sup>o</sup> | .356 |
| **SSRT – P3 pl** | .029 | .277 | .150 | .357 | .140 | .229 | .079 | .162 | .035 |
| **SSRT – EMG** | .390 | .645 | .573 | .669 | .360 | .447 | .682<sup>o</sup> | .737 | .720 |
| **EMG – P3 ol** | .615 | .420 | .561 | .752 | .437<sup>o</sup> | .562<sup>o</sup> | -.115 | .060<sup>o</sup> | .540 |
| **EMG – P3 pl** | .562 | -.086 | .328 | .437 | -.083 | .371<sup>o</sup> | -.105 | .305 | .106 |
| **GoRT – P3 ol** | .077 | -.093 | .001 | -.059 | -.395 | -.275 | -.278 | -.279 | -.256 |
| **GoRT – P3 pl** | .009 | .253 | .120 | -.171 | -.547 | -.235 | -.226 | -.418 | .010 |
| **GoRT – EMG** | -.356 | -.457 | -.474 | -.027 | -.466 | -.485 | -.118 | -.403 | -.427 |
| **GoRT – SSRT** | -.283 | -.608 | -.269 | -.056 | -.422 | -.331 | -.312 | -.156 | -.169 |

<sup>o</sup> indicates correlations where a bivariate outlier was removed (N = 17)
Abbreviations: pl – peak latency, ol – onset latency, SSRT – stop signal reaction time, GoRT – go reaction time

The SSRT and P3 onset latency showed medium to large positive correlations at almost all measurement points (BF_10_ = 12.721, *M*_*r*_ = .418), suggesting that faster inhibition as measured by the SSRT was accompanied by earlier onsets of the P3 potential. The relationship between the SSRT and P3 peak latency was also generally positive (BF_10_ = 19.042, *M*_*r*_ = .164). Furthermore, earlier prEMG latencies were related to shorter SSRTs (BF_10_ = 1706.279, *M*_*r*_ = .595) as well as earlier onsets of the P3 (BF_10_ = 21.029, *M_r_ =* .460). The correlations between prEMG latencies and P3 peak latencies were more variable both in size and direction (BF_10_ = 2.110, *M_r_ =* .220).

To assess the relationship between inhibition measures and performance in go trials, we calculated correlation coefficients between goRT and P3 peak and onset latencies, SSRTs, and prEMG peak latencies. Longer reaction times were generally associated with shorter latencies for the P3 onset (BF_10_ = 5.636, *M_r_ =* -.177), shorter prEMG peak latency (BF_10_ = 131.931, *M_r_ =* -.365), as well as shorter SSRTs (BF_10_ = 35.060, *M_r_ =* -.299), while the relationship between goRTs and P3 peak latencies showed more variability across measurements (BF_10_ = 0.810, *M_r_ =* -.144).

### tDCS questionnaire

We also investigated the presence and severity of potential side effects of tDCS using a questionnaire suggested by Brunoni et al^41^. Participants reported few side effects of tDCS, with itching and tingling being reported most commonly. The reported severity of these effects did not differ between conditions (BF_10_ = 0.277). Participants were also asked to rate whether they thought the experienced side effects were related to tDCS. The data was inconclusive regarding whether participants were more likely to attribute side effects to active stimulation compared to SHAM (BF_10_ = 1.126), potentially due to the low amount of side effects reported in general.

## Discussion

To assess the contribution of sensory and inhibitory processing to the latency of stopping, we investigated the effects of tDCS over the IFG and VC on different putative measures of inhibitory timing, namely the SSRT, P3 onset and peak latencies, and the latency of the prEMG. Since several different markers have been associated with the timing of inhibition, we also assessed the relationship between these markers, as well as their relationship with go trial reaction times.

Contrary to our expectations, we found evidence against any stimulation-specific effects on any of the investigated behavioral measures for both stimulation conditions, thus showing that tDCS did not affect reaction times in go trials, SSDs or SSRTs. The evidence also suggested that tDCS did not affect P3 latency measures or prEMG latencies. Therefore, our results contrast with previous findings of SSRT reductions following anodal IFG stimulation (but see ^10^ for an exception). This discrepancy could be driven by several methodological differences between this and earlier reports, such as experimental design, electrode locations, or stimulation lateralization.

First, we utilized a within-subject design with several active and sham stimulation conditions in combination with a multimodal approach and pre-, peri- and post-stimulation measurements. This allowed for a more thorough examination of potential tDCS effects on response inhibition than what has been possible in previous studies. As such, our evidence for H0 shows that additional control conditions are necessary when researching the effects of tDCS on stopping.

Second, whereas previous studies have largely relied on approximate and thus unprecise electrode positions to stimulate the IFG, we chose electrode positions after modelling the current distribution for a given target region. Consequently, the discrepant results might be caused by differences in electrode positioning, both with respect to the exact position of the anode as well as the resulting current distributions. Previous SSRT decreases have been reported after anodal stimulation of the right IFG only, usually with a contralateral and often more anterior cathode. However, these types of distant bipolar montages can result in widespread and diffuse neuromodulation^32,42,43^, which raises the possibility that other regions might drive the previously reported effects. Furthermore, while high-definition tDCS to the right IFG has been associated with an SSRT decrease (albeit with smaller effects than conventional montages)^32^, several studies point to the involvement of bilateral IFG in SST performance^44,45^. For these reasons, we used an electrode montage that provided simultaneous anodal stimulation of the bilateral IFG with ipsilateral cathodes placed posterior to the anodes. This setup provided more control over effects that might stem from the accidental stimulation of larger prefrontal and/or premotor areas. We also chose the same cathodal placement for both active stimulation conditions for better control over potential cathodal effects. While the decreased distance between anode and cathode has been found to provide more focal stimulation, it comes at the cost of higher amounts of current being shunted across the scalp^43^. Therefore, we stimulated at a higher intensity compared to previous studies investigating tDCS effects on stopping. In sum, our results show that further investigations are needed to ascertain whether SSRT reductions following anodal IFG stimulation are actually driven by the concurrent modulation of neural activity outside of the IFG.

Concerning the relationship between the different proposed markers of stopping latency, our results were in accordance with previous studies. We found positive correlations between the P3 onset latencies and SSRTs^13–15^, as well as between prEMG latencies and SSRTs^17^. In addition, we also found positive correlations between prEMG and P3 onset latencies. In sum, this suggests that earlier EMG activity is followed by earlier SSRTs, and an earlier onset of the P3. Furthermore, the correlations between these measures and reaction times in go trials were negative, in line with the notion that proactive slowing of behavioral responses can facilitate inhibitory control^46^. However, the markers differ noticeably in their estimated stopping latency (Figure 2). Our EMG results suggests that the decline of motor activity occurred at around 160 ms after stop-signal presentation, thus replicating previous findings regarding the timing of motor suppression in response effectors^17,18^, and mirroring the timing of inhibition measured at the cortical level^47^. This is considerably earlier than the latency of the SSRT and the P3 onset, which in this study were estimated to be 233 ms and 302 ms, respectively. Not least, even with correlations as high as. 59, as seen between the SSRT and the prEMG, the common explained variance is only about 35%. While this corresponds to a large effect, it leaves much unexplained variance that could be driven by factors unique to each measure. Therefore, it seems reasonable to question the extent to which these markers should be used interchangeably as measures of inhibitory timing, a notion which is supported by recent work^48,49^. In sum, though, the correlational pattern suggests that while the different proposed stopping latency markers are not necessarily overlapping in time (Figure 2), they are indeed related to each other as well as to general task processing as quantified by the go reaction times.

**Figure 2.**
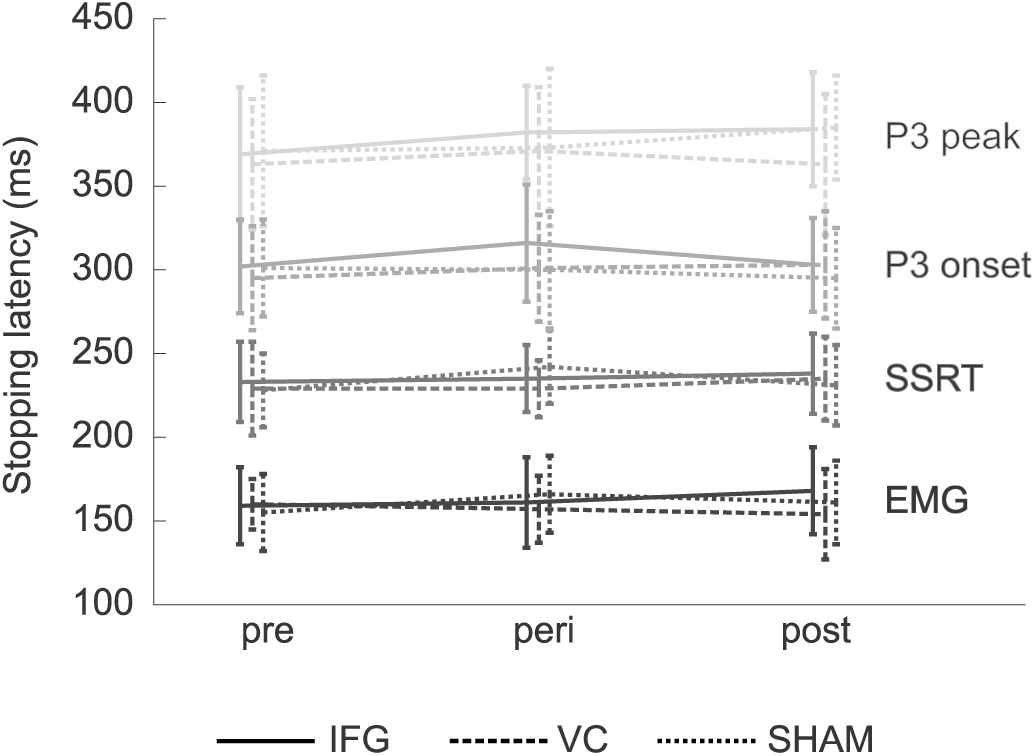
Mean latency estimates derived from the different putative measures of stopping latency, estimated separately for each measurement. Error bars represent *SD*.

Two potential limitations to the tDCS results necessitates further discussion. First, since we used stimulation electrodes placed over the visual cortex, we were not able to simultaneously record EEG data over occipital areas, thus prohibiting any investigation of how the different stimulation protocols could have affected ERP components associated with early sensory processing. Second, we found a general condition effect on reaction times, with faster goRTs in the IFG condition compared to the VC condition. Importantly, the RT effects were accompanied by similar differences in SSDs, thus yielding comparable SSRT estimates between conditions. One potential explanation for this effect is that the position of the stimulation electrodes affected participants’ expectations and subsequent behavior. Since electrodes were positioned prior to the first run of the task, this effect could occur already pre-stimulation. Due to balancing sham electrode placements between the two active conditions, the resulting sample size for each sham setup was too small to properly investigate this possibility. However, this has important implications for the selection of electrode sites for active control conditions, and suggests that the investigation of moderating effects, including those related to participants’ expectancies, need to be more stringently integrated into experimental designs.

## Conclusion

This study shows that neither tDCS to the IFG nor to the VC modulated any of the suggested markers of stopping latency, which challenges previous studies arguing that a single session of anodal tDCS over the IFG can improve inhibition. As such, it illustrates the need for more research utilizing several control conditions and concurrent neural measures to establish potential effects of tDCS on inhibitory control. Our results also suggest that while EMG latencies, SSRTs and P3 onsets show a positive relationship with each other, they provide different estimates for the timing of inhibition. This suggests that using these markers interchangeably as measures of inhibitory capabilities is premature, and that future studies should aim to further disentangle what is reflected in the different signatures associated with stopping.

## Methods

### Participants

28 healthy participants were recruited for the study. However, 4 participants failed to complete all three measurements, and 6 additional participants were excluded from further analyses. Of those 6, two were excluded for having longer RTs in unsuccessful stop trials compared to go trials, thus violating the assumptions of the independent horse race model. Three participants were excluded because their stopping accuracy was too low to ensure an acceptable signal-to-noise ratio in the ERPs. An additional participant was excluded because behavioral results indicated that task instructions were not followed correctly. This resulted in 18 participants (9 females, mean age = 24 years) for further analysis. All participants were naïve to tDCS and reported no contraindications for electrical stimulation. In accordance with the Helsinki declaration, all participants provided written consent prior to participation, and everyone received monetary compensation. The study was approved by the institutional review board of the Department of Psychology, University of Oslo.

### Task and procedure

Each tDCS-session consisted of three runs of two experimental tasks (pre-, peri- and post-stimulation) with concurrent EEG- and EMG-recordings (Figure 3a), and a tDCS-related questionnaire^41^. The tasks were a visual SST and a two-choice reaction time (CRT) task presented in alternating blocks. For this report, we focused exclusively on the SST (Figure 3b). Each task run consisted of 450 SST trials with a stop-signal probability of. 24. Colored arrows were used as stimuli, where the direction of the arrow indicated the responding hand. All stimuli were centrally presented against a grey background, and stimulus color assignments (green, orange and blue for CRT-go, SST-go and SST-stop stimuli) were counterbalanced across participants. The stop-signal delay (SSD) was adjusted based on a tracking algorithm which increased or decreased the SSD by 50 ms following successful and unsuccessful stop trials, respectively. Performance feedback was given after every 75^th^ trial, and participants were instructed to be faster if the average goRT was above 600 ms, and to be more accurate if the average stop accuracy was below. 40.

**Figure 3.**
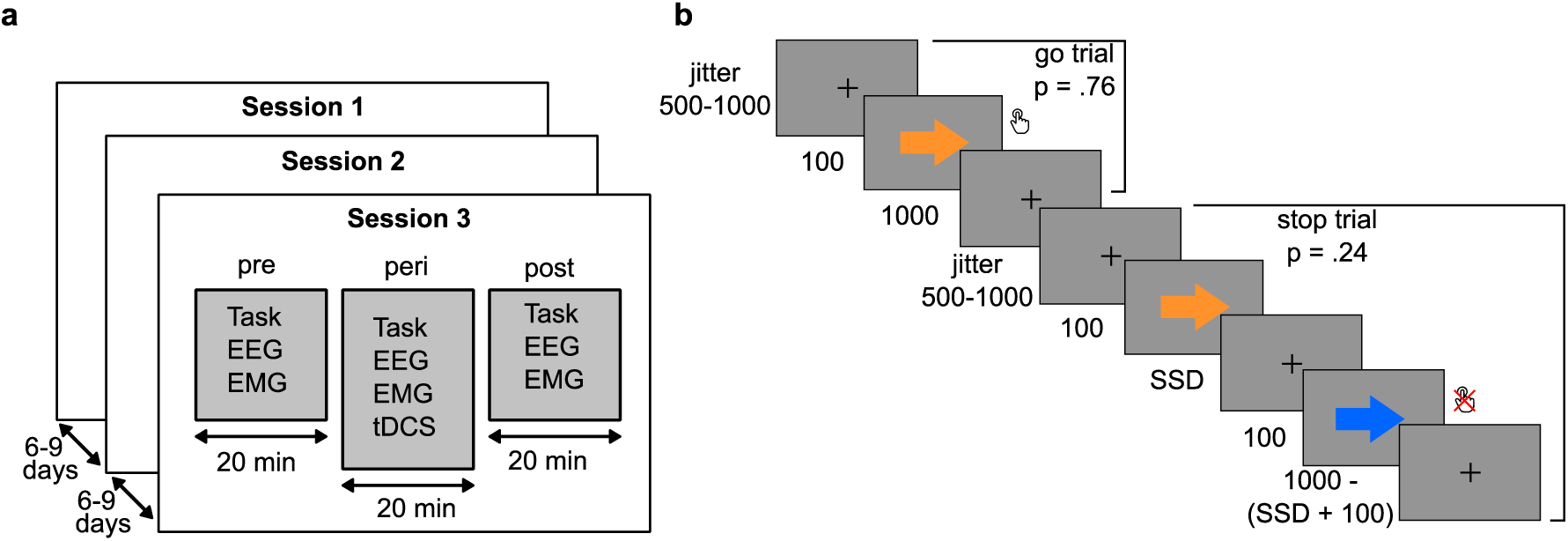
**a.** All participants completed three separate tDCS sessions, each session consisting of pre-, peri- and post-stimulation measurements. **b.** Overview of the stop-signal task with the timing of the stimuli given in ms.

### Data acquisition

Both EEG- and EMG-activity were recorded using a BrainAmp system (Brain Products GmbH, Germany) (Figure 4a and 4b). We used online low-pass filters at 250 Hz (EEG) and 1000 Hz (EMG), a sampling rate of 5000 Hz, and a resolution of 0.5 µV. EEG electrode impedances during recording were kept equal to or below 5 kΩ. For tDCS, all participants received tDCS over the IFG, the visual cortex (VC) and sham stimulation (Figure 4c and 4d). Participants were blind to the different conditions, and condition order was randomized. TDCS was given using a NeuroConn DC-stimulator MC (GmbH, Germany), using a multifocal setup with two 5×5 cm rubber electrodes on each hemisphere. TDCS electrode impedances were kept equal to or below 10 kΩ.

**Figure 4.**
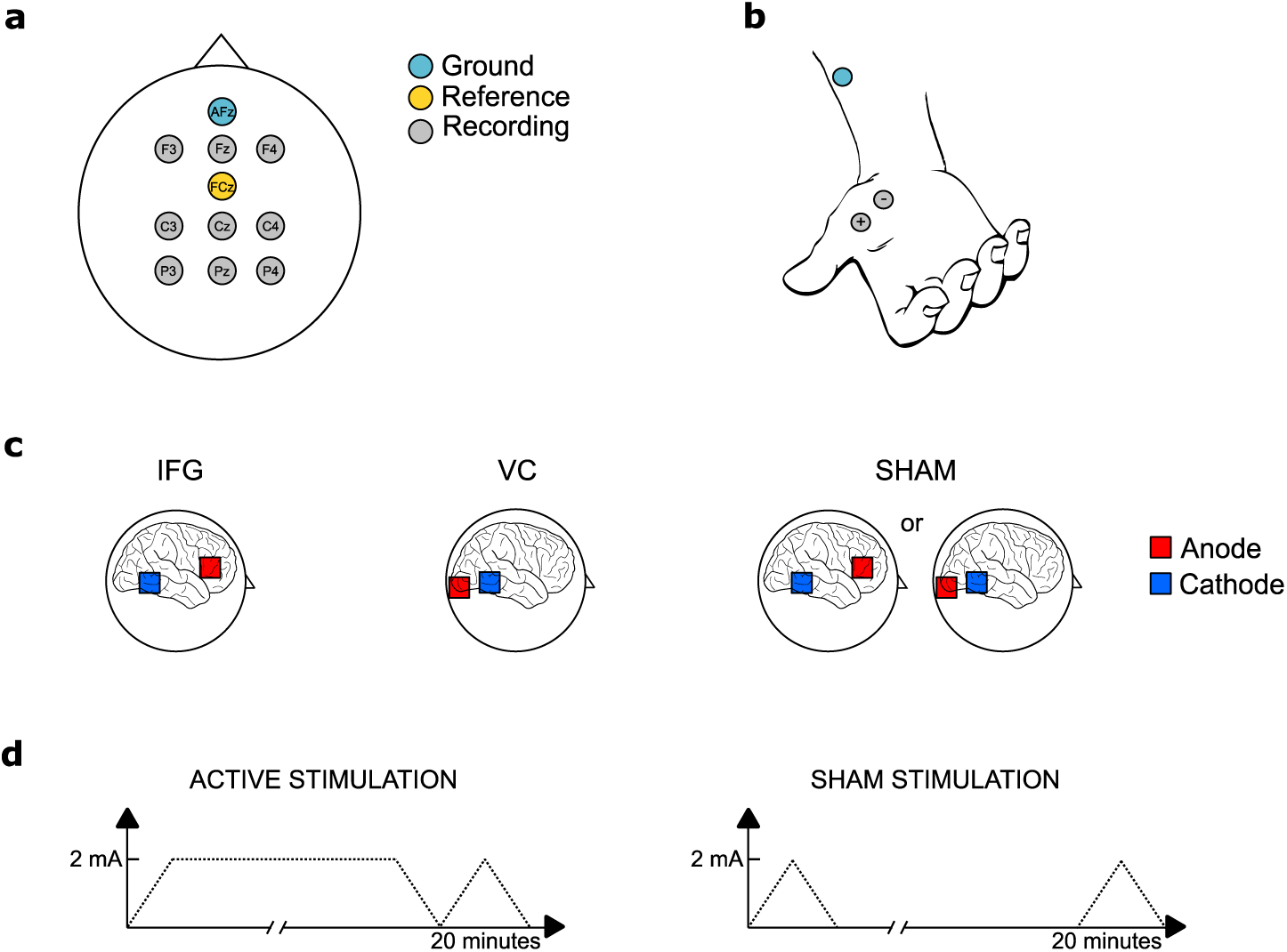
**a.** EEG activity was recorded from 9 electrodes positioned at F3, Fz, F4, C3, Cz, C4, P3, Pz, P4 according to the international 10-20 system. The online reference and ground electrode were placed at FCz and AFz, respectively. Additional electrodes were positioned at both earlobes and the tip of the nose for offline re-referencing. **b.** EMG activity was recorded from the *abductor pollicis brevis* on each hand using a bipolar electrode montage. Ground electrodes were placed on each forearm. **c.** Anodal electrodes (red) were centered at FC7 and FC8 (IFG-condition) or O1 and O2 (VC-condition). Cathodal placement (blue) was identical for both conditions, with the top of the electrode aligned with CP8 and P8 and CP7 and P7. For the SHAM session, half of the participants received the IFG-setup, while the other half received the VC-setup. **d.** In the active stimulation sessions, tDCS was ramped up to 2mA over 30 seconds, applied for 20 minutes, and then ramped down. The SHAM session only included the ramps.

### Preprocessing

EEG preprocessing consisted of visual identification and exclusion of time frames containing tDCS-related artifacts, re-referencing to averaged earlobes, low-pass-filtering (40 Hz), down-sampling (500 Hz), high-pass-filtering (0.1 Hz), ocular artifact correction based on independent component analysis, epoching (−200ms to +800ms relative to stop-stimulus onset), baseline correction and threshold based artifact rejection (absolute threshold = 120 µV). EMG preprocessing consisted of low-pass-filtering (250 Hz), down-sampling (500 Hz) and high-pass-filtering (20 Hz). All filtering was performed using the basic FIR filter implemented in EEGLAB^50^. Then, EMG time courses were extracted for all trial types (−200ms to +1600ms relative to go-stimulus onset) and baseline corrected. The EMG activity was transformed and smoothed in the time-domain (for details, see ^17^) and z-scored across all trials. The resulting EMG time courses reflect the ratio of a root mean square-transformed signal relative to baseline, z-scored relative to the total amount of EMG activity in all trials. Following this, trials where the average untransformed baseline activity exceeded 100 µV were defined as artifacts and excluded from further analyses. Due to electrical noise of the DC stimulator and for baseline estimation purposes only, the data was notch-filtered at 100 Hz and 200 Hz using a basic FIR filter with a −6 dB roll-off rate and passband edges of 95/105 Hz and 195/205 Hz, respectively. Following artifact rejection, an automated algorithm was used to detect the presence of EMG activity in successful stop trials. Specifically, an EMG response was detected if any time point between go-stimulus onset and the response window offset exceeded 1.2, i.e. activity that deviated from baseline with at least 1.2 SDs. The SD threshold was based on visual examination of the data, and detection algorithm performance was visually validated in a random subsample of trials. After this, the time courses were re-epoched (−200ms to +800 ms relative to stop-stimulus onset), baseline corrected, and averaged across trials.

### Statistical analyses

From the EEG, we extracted peak and onset latencies of the stop-P3 potential at electrode Cz. Peak latency was defined as the latency of the local maximum within 250 – 500 ms post-stop, while P3 onset latency was defined as the half-amplitude latency, i.e. the earliest latency at which the P3 exceeded 50 % of its peak amplitude when tracking amplitudes backwards in time from the peak. This approach has been found to be a reliable onset estimation procedure for late potentials like the P3, with acceptable power for detecting relatively small differences in onset latency^51^. From the EMG, we extracted the percentage of detected prEMG responses from successful stop trials, as well as the peak latency of these responses. Peak latency was extracted from the averaged prEMG time courses and defined as the maximum between stop-signal onset and the participant’s SSRT for that session and measurement. From the behavioral data, we extracted reaction times in go trials (goRTs), reaction times in unsuccessful stop trials (USRTs), SSRTs (using the integration method^52^) and average stop-signal delays (SSDs).

All statistical analyses were performed within a Bayesian framework using the JASP software^37^, and Bayes factors (BF_10_) are reported. BFs express the relative likelihood of the data under two competing hypotheses, essentially providing an estimate of how likely the alternative hypothesis is compared to the null hypothesis^38^.

First, the data was assessed for outliers, defined as values with a z-score of +/-3.5. None of the participants qualified as outliers on any of the analyzed measures. Then, to ensure the validity of the horse-race model, individual paired-samples t-tests were performed comparing goRTs to USRTs for each condition and time point using a default Cauchy (0, r = 0.707) prior.

Behavioral, EEG and EMG data were analyzed using two-way Bayesian repeated measures ANOVAs^53^ (rmANOVA), with the factors **time** (pre, peri, post) and **condition** (IFG, VC, SHAM). We used default priors, i.e. a multivariate Cauchy (r scale fixed effects = 0.5, r scale random effects = 1, r scale covariates = 0.354) distribution^54^, and set number of samples to 100 000 for precise BF estimation. BFs for main effects indicating at least moderate evidence in favor of the alternative hypothesis were followed up with post hoc tests, which are based on t-tests with a default Cauchy (0, r = 0.707) prior. Sphericity was assessed using Mauchly’s *W*, and Greenhouse-Geisser estimated epsilon (ε) will be provided. However, as correction procedures for sphericity violations are not yet well developed within the Bayesian framework, no correction could be implemented.

To investigate the relationship between the different inhibition indices, we calculated pairwise correlations between SSRTs, prEMG peak latency, P3 peak latency and P3 onset latency separately for each condition and time point. Furthermore, to explore the relationship between inhibition measures and go trial performance, we calculated correlations between goRT and P3 peak and onset latencies, as well as between goRT and the prEMG latency. Bivariate outliers, defined as highly influential values quantified by a Cook’s *d* exceeding 1^55^, were removed prior to analysis. To get an indication of the stability of the direction and size the obtained correlations, the correlation coefficients were Fisher-z-transformed^56^ and tested against the null hypothesis with a one-sample t-test using a default Cauchy (0, r = 0.707) prior.

Lastly, as an evaluation of tDCS safety and condition blinding, we analyzed responses to a tDCS questionnaire^41^. Participants indicated the presence of a range of potential side effects using a scale from 1-4 (PRESENCE), as well as if they thought any experienced effects were caused by the stimulation using a scale from 1-5 (PROBABILITY). Estimates were obtained for each participant by averaging across their presence and probability scores separately. Presence and probability scores were analyzed using separate rmANOVAs with **condition** (IFG, VC and SHAM) as a factor and the default Cauchy priors.

## Data availability

The datasets generated and analyzed as part of the current study are available from the corresponding author on reasonable request.

## Author Contributions Statement

CNT, MSM, LR and RJH conceived and planned the study. CNT and MSM carried out the experiment. CNT performed the analyses under the guidance of RJH and LR. CNT wrote the manuscript. All authors edited the manuscript and agreed to the final version.

## Additional information

The authors declare no competing interests.

## References

1 Aron, A. R., Robbins, T. W. & Poldrack, R. A. Inhibition and the right inferior frontal cortex. Trends Cogn Sci 8, 170–177, doi:10.1016/j.tics.2004.02.010 (2004).

2 Aron, A. R., Robbins, T. W. & Poldrack, R. A. Inhibition and the right inferior frontal cortex: one decade on. Trends Cogn Sci 18, 177–185, doi:10.1016/j.tics.2013.12.003 (2014).

3 Logan, G. D. & Cowan, W. B. On the Ability to Inhibit Thought and Action - a Theory of an Act of Control. Psychological review 91, 295–327, doi:10.1037/0033-295x.91.3.295 (1984).

4 de Jong, R., Coles, M. G. H., Logan, G. D. & Gratton, G. In search of the point of no return: the control of response processes. Journal of experimental psychology. Human perception and performance 16, 164–182, doi:10.1037/0096-1523.16.1.164 (1990).

5 Enriquez-Geppert, S., Konrad, C., Pantev, C. & Huster, R. J. Conflict and inhibition differentially affect the N200/P300 complex in a combined go/nogo and stop-signal task. Neuroimage 51, 877–887, doi:10.1016/j.neuroimage.2010.02.043 (2010).

6 Greenhouse, I. & Wessel, J. R. EEG signatures associated with stopping are sensitive to preparation. Psychophysiology 50, 900–908, doi:10.1111/psyp.12070 (2013).

7 Hoptman, M. J. et al. Sensory and cross-network contributions to response inhibition in patients with schizophrenia. NeuroImage: Clinical 18, 31–39, doi:10.1016/j.nicl.2018.01.001 (2018).

8 Johnstone, S. J. et al. The development of stop-signal and Go/Nogo response inhibition in children aged 7–12 years: Performance and event-related potential indices. International Journal of Psychophysiology 63, 25–38, doi:10.1016/j.ijpsycho.2006.07.001 (2007).

9 Logemann, H. N., Bocker, K. B., Deschamps, P. K., Kemner, C. & Kenemans, J. L. The effect of enhancing cholinergic neurotransmission by nicotine on EEG indices of inhibition in the human brain. Pharmacol Biochem Behav 122, 89–96, doi:10.1016/j.pbb.2014.03.019 (2014).

10 Cunillera, T., Brignani, D., Cucurell, D., Fuentemilla, L. & Miniussi, C. The right inferior frontal cortex in response inhibition: A tDCS-ERP co-registration study. Neuroimage 140, 66–75, doi:10.1016/j.neuroimage.2015.11.044 (2016).

11 Palmwood, E. N., Krompinger, J. W. & Simons, R. F. Electrophysiological indicators of inhibitory control deficits in depression. Biol Psychol 130, 1–10, doi:10.1016/j.biopsycho.2017.10.001 (2017).

12 Kusztor, A. et al. Sleep deprivation differentially affects subcomponents of cognitive control. Sleep 42, doi:10.1093/sleep/zsz016 (2019).

13 Wessel, J. R. Prepotent motor activity and inhibitory control demands in different variants of the go/no-go paradigm. Psychophysiology 55, doi:10.1111/psyp.12871 (2018).

14 Wessel, J. R. & Aron, A. R. It’s not too late: the onset of the frontocentral P3 indexes successful response inhibition in the stop-signal paradigm. Psychophysiology 52, 472–480, doi:10.1111/psyp.12374 (2015).

15 Wessel, J. R. et al. Surprise disrupts cognition via a fronto-basal ganglia suppressive mechanism. Nat Commun 7, 11195, doi:10.1038/ncomms11195 (2016).

16 Anguera, J. A. & Gazzaley, A. Dissociation of motor and sensory inhibition processes in normal aging. Clinical Neurophysiology 123, 730–740, doi:10.1016/j.clinph.2011.08.024 (2012).

17 Raud, L. & Huster, R. J. The Temporal Dynamics of Response Inhibition and their Modulation by Cognitive Control. Brain Topogr 30, 486–501, doi:10.1007/s10548-017-0566-y (2017).

18 Raud, L., Westerhausen, R., Dooley, N. & Huster, R. J. Differences in unity: the go/no-go and stop signal tasks rely on different inhibitory mechanisms. bioRxiv, 705079, doi:10.1101/705079 (2019).

19 Logan, G. D., Van Zandt, T., Verbruggen, F. & Wagenmakers, E. J. On the ability to inhibit thought and action: general and special theories of an act of control. Psychological review 121, 66–95, doi:10.1037/a0035230 (2014).

20 Verbruggen, F., McLaren, I. P. & Chambers, C. D. Banishing the Control Homunculi in Studies of Action Control and Behavior Change. Perspect Psychol Sci 9, 497–524, doi:10.1177/1745691614526414 (2014).

21 Verbruggen, F., Stevens, T. & Chambers, C. D. Proactive and reactive stopping when distracted: an attentional account. Journal of experimental psychology. Human perception and performance 40, 1295–1300, doi:10.1037/a0036542 (2014).

22 Boucher, L., Palmeri, T. J., Logan, G. D. & Schall, J. D. Inhibitory control in mind and brain: an interactive race model of countermanding saccades. Psychological review 114, 376–397, doi:10.1037/0033-295X.114.2.376 (2007).

23 Salinas, E. & Stanford, T. R. The countermanding task revisited: fast stimulus detection is a key determinant of psychophysical performance. The Journal of neuroscience: the official journal of the Society for Neuroscience 33, 5668–5685, doi:10.1523/JNEUROSCI.3977-12.2013 (2013).

24 Jahfari, S., Ridderinkhof, K. R. & Scholte, H. S. Spatial frequency information modulates response inhibition and decision-making processes. PLoS One 8, e76467, doi:10.1371/journal.pone.0076467 (2013).

25 Boehler, C. N. et al. Sensory MEG responses predict successful and failed inhibition in a stop-signal task. Cereb Cortex 19, 134–145, doi:10.1093/cercor/bhn063 (2009).

26 Langford, Z. D., Krebs, R. M., Talsma, D., Woldorff, M. G. & Boehler, C. N. Strategic down-regulation of attentional resources as a mechanism of proactive response inhibition. Eur J Neurosci 44, 2095–2103, doi:10.1111/ejn.13303 (2016).

27 Montanari, R., Giamundo, M., Brunamonti, E., Ferraina, S. & Pani, P. Visual salience of the stop-signal affects movement suppression process. Exp Brain Res 235, 2203–2214, doi:10.1007/s00221-017-4961-0 (2017).

28 Nitsche, M. A. & Paulus, W. Excitability changes induced in the human motor cortex by weak transcranial direct current stimulation. J Physiol 527 Pt 3, 633–639 (2000).

29 Cai, Y. et al. The Role of the Frontal and Parietal Cortex in Proactive and Reactive Inhibitory Control: A Transcranial Direct Current Stimulation Study. Journal of Cognitive Neuroscience 28, 177–186, doi:10.1162/jocn_a_00888 (2015).

30 Castro-Meneses, L. J., Johnson, B. W. & Sowman, P. F. Vocal response inhibition is enhanced by anodal tDCS over the right prefrontal cortex. Exp Brain Res 234, 185–195, doi:10.1007/s00221-015-4452-0 (2016).

31 Cunillera, T., Fuentemilla, L., Brignani, D., Cucurell, D. & Miniussi, C. A simultaneous modulation of reactive and proactive inhibition processes by anodal tDCS on the right inferior frontal cortex. PLoS One 9, e113537, doi:10.1371/journal.pone.0113537 (2014).

32 Hogeveen, J. et al. Effects of High-Definition and Conventional tDCS on Response Inhibition. Brain Stimul 9, 720–729, doi:10.1016/j.brs.2016.04.015 (2016).

33 Jacobson, L., Javitt, D. C. & Lavidor, M. Activation of Inhibition: Diminishing Impulsive Behavior by Direct Current Stimulation over the Inferior Frontal Gyrus. Journal of Cognitive Neuroscience 23, 3380–3387, doi:10.1162/jocn_a_00020 (2011).

34 Stramaccia, D. F. et al. Assessing the effects of tDCS over a delayed response inhibition task by targeting the right inferior frontal gyrus and right dorsolateral prefrontal cortex. Exp Brain Res 233, 2283–2290, doi:10.1007/s00221-015-4297-6 (2015).

35 Ditye, T., Jacobson, L., Walsh, V. & Lavidor, M. Modulating behavioral inhibition by tDCS combined with cognitive training. Exp Brain Res 219, 363–368, doi:10.1007/s00221-012-3098-4 (2012).

36 Lee, C., Jung, Y.-J., Lee, S. J. & Im, C.-H. COMETS2: an advanced MATLAB toolbox for the numerical analysis of electric fields generated by transcranial direct current stimulation. Journal of neuroscience methods 277, 56–62 (2017).

37 37 JASP (Version 0.9) (2018).

38 Wagenmakers, E.-J. et al. Bayesian inference for psychology. Part I: Theoretical advantages and practical ramifications. Psychonomic Bulletin & Review 25, 35–57, doi:10.3758/s13423-017-1343-3 (2018).

39 Jeffreys, H. The theory of probability. (OUP Oxford, 1998).

40 Lee, M. D. & Wagenmakers, E.-J.. Bayesian cognitive modeling: A practical course. (Cambridge university press, 2014).

41 Brunoni, A. R. et al. A systematic review on reporting and assessment of adverse effects associated with transcranial direct current stimulation. Int J Neuropsychopharmacol 14, 1133–1145, doi:10.1017/S1461145710001690 (2011).

42 Csifcsák, G., Boayue, N. M., Puonti, O., Thielscher, A. & Mittner, M. Effects of transcranial direct current stimulation for treating depression: A modeling study. Journal of affective disorders 234, 164–173 (2018).

43 Datta, A., Elwassif, M., Battaglia, F. & Bikson, M. Transcranial current stimulation focality using disc and ring electrode configurations: FEM analysis. Journal of neural engineering 5, 163 (2008).

44 Swick, D., Ashley, V. & Turken, A. U. Left inferior frontal gyrus is critical for response inhibition. BMC Neurosci 9, 102, doi:10.1186/1471-2202-9-102 (2008).

45 Li, C. S., Huang, C., Constable, R. T. & Sinha, R. Imaging response inhibition in a stop-signal task: neural correlates independent of signal monitoring and post-response processing. The Journal of neuroscience: the official journal of the Society for Neuroscience 26, 186–192, doi:10.1523/jneurosci.3741-05.2006 (2006).

46 Aron, A. R. From reactive to proactive and selective control: developing a richer model for stopping inappropriate responses. Biol Psychiatry 69, e55–68, doi:10.1016/j.biopsych.2010.07.024 (2011).

47 van den Wildenberg, W. P. M. et al. Mechanisms and Dynamics of Cortical Motor Inhibition in the Stop-signal Paradigm: A TMS Study. Journal of Cognitive Neuroscience 22, 225–239, doi:10.1162/jocn.2009.21248 (2010).

48 Huster, R. J., Messel, M. S., Thunberg, C. & Raud, L. The P300 as marker of inhibitory control – fact or fiction. bioRxiv, 694216, doi:10.1101/694216 (2019).

49 Skippen, P. et al. Reconsidering electrophysiological markers of response inhibition in light of trigger failures in the stop-signal task. bioRxiv, 658336, doi:10.1101/658336 (2019).

50 Delorme, A. & Makeig, S. EEGLAB: an open source toolbox for analysis of single-trial EEG dynamics including independent component analysis. Journal of Neuroscience Methods 134, 9–21, doi:10.1016/j.jneumeth.2003.10.009 (2004).

51 Kiesel, A., Miller, J., Jolicœur, P. & Brisson, B. Measurement of ERP latency differences: A comparison of single‐participant and jackknife‐based scoring methods. Psychophysiology 45, 250–274 (2008).

52 Verbruggen, F., Chambers, C. D. & Logan, G. D. Fictitious inhibitory differences: how skewness and slowing distort the estimation of stopping latencies. Psychol Sci 24, 352–362, doi:10.1177/0956797612457390 (2013).

53 BayesFactor (Version 0.9.11-3) (2015).

54 Rouder, J. N., Morey, R. D., Speckman, P. L. & Province, J. M. Default Bayes factors for ANOVA designs. Journal of Mathematical Psychology 56, 356–374 (2012).

55 Cook, R. D. & Weisberg, S. Residuals and influence in regression. (New York: Chapman and Hall, 1982).

56 Fisher, R. A. Frequency distribution of the values of the correlation coefficient in samples from an indefinitely large population. Biometrika 10, 507–521 (1915).

